# Discovery of RNA-Protein Molecular Clamps Using Proteome-Wide Stability Assays

**DOI:** 10.1101/2024.04.19.590252

**Authors:** Stanley I. Goldstein, Alice C. Fan, Zihao Wang, Sai K. Naineni, Regina Cencic, Steve B. Garcia-Gutierrez, Kesha Patel, Sidong Huang, Lauren E. Brown, Andrew Emili, John A. Porco

## Abstract

Uncompetitive inhibition is an effective strategy for suppressing dysregulated enzymes and their substrates, but discovery of suitable ligands depends on often-unavailable structural knowledge and serendipity. Hence, despite surging interest in mass spectrometry-based target identification, proteomic studies of substrate-dependent target engagement remain sparse. Herein, we describe a strategy for the discovery of substrate-dependent ligand binding. Using proteome integral solubility alteration (PISA) assays, we show that simple biochemical additives can enable detection of RNA-protein-small molecule complexes in native cell lysates. We apply our approach to rocaglates, molecules that specifically clamp RNA to eukaryotic translation initiation factor 4A (eIF4A), DEAD-box helicase 3X (DDX3X), and potentially other members of the DEAD-box (DDX) helicase family. To identify unexpected interactions, we used a target class-specific thermal window and compared ATP analog and RNA base dependencies for key rocaglate-DDX interactions. We report and validate novel DDX targets of high-profile rocaglates – including the clinical candidate Zotatifin – using limited proteolysis-mass spectrometry and fluorescence polarization (FP) experiments. We also provide structural insight into divergent DDX3X affinities between synthetic rocaglates. Taken together, our study provides a model for screening uncompetitive inhibitors using a chemical proteomics approach and uncovers actionable DDX clamping targets, clearing a path towards characterization of novel molecular clamps and associated RNA helicases.

## INTRODUCTION

Rocaglates are a diverse family of natural products that have been subject to numerous synthetic and medicinal chemistry campaigns across academia and industry.^1–13^ Initial interest in their pharmacology arose from studies of crude *Aglaia* extracts showing anti-neoplastic activity in cellular and murine models of cancer.^14^ After the prototypical rocaglates RocA and silvestrol were found to drive these effects^15,16^, investigations in models of viral, protozoan, and bacterial infection highlighted their broader utility in infectious disease.^17–25^ Our group and others have since expanded the rocaglate family to hundreds of derivatives and novel subclasses, each with demonstrated activity across a range of disease models^3,6,12,26^ – representative structures of rocaglates used in this study are provided in **Figure S1A**. Sustained interest in rocaglate drug discovery has culminated in the nomination of Zotatifin (eFT226) as a clinical candidate for both estrogen receptor-positive (ER+) breast cancer and SARS-CoV-2 infection (ClinicalTrials.gov identifiers: NCT04092673 and NCT04632381), underscoring the tractability of rocaglates as broad-spectrum pharmaceutical agents.

The bulk of mechanistic studies converge on cap-dependent translation as a nexus for rocaglate bioactivity.^27–31^ Early experiments established rocaglates as translation inhibitors that stabilize the association between mRNA and eukaryotic translation initiation factor 4A1 (eIF4A1)^32,33^, a DEAD-box helicase (DDX) that unwinds secondary structure in transcripts destined for translation. This stabilization, hereby referred to as “clamping,” halts the dissociation of eIF4A1 from purine-rich 5’-UTRs, resulting in the steric blockade of ribosome recruitment.^31^ A crystallographic study by Iwasaki and coworkers revealed that rocaglates bind in a transient cleft between eIF4A1 and purine dinucleotides, thereby forming contacts with both biomolecules.^34^ It was later shown by the Pelletier laboratory that rocaglates clamp eIF4A2, a paralog of eIF4A1 also implicated in translation^26^, and eIF4A3 (also known as DDX48), a paralog involved in exon-junction complex (EJC) formation, although the consequences of eIF4A3 clamping have yet to be fully characterized.^30^ Finally, a mass spectrometry study by Chen and coworkers identified the DEAD-box helicase 3X (DDX3X) – another protein involved in translation initiation, albeit with a less well-defined role – as a low affinity target of RocA.^29^

The aforementioned four helicases share a highly conserved core domain common to at least 41 human DDX and 16 related DExH-box (DHX, hereafter binned with DDX for simplicity) helicases whose functions have been reviewed in detail.^35,36^ The core domain provides ATPase and RNA binding activity, while the variable N- and C-terminal domains give rise to diverse functions across RNA-related processes including translation, rRNA processing, and antiviral innate immunity.^37–40^ Given their broad roles, DDXs represent an attractive class of drug targets if they prove to be selectively ligandable. Thus, the understanding that eIF4A and DDX3X belong to a larger family of DDXs invites speculation as to whether synthetic rocaglates can be engineered to selectively target other DDX proteins. However, undifferentiated targeting of eIF4A paralogs and DDX3X clouds distinctions between biological effects arising downstream of ligand binding.^28–30^ Selectivity becomes relevant in specific disease subtypes that may preferentially depend on certain helicases. As such, the identification of rocaglates selective for specific DDX targets remains a key challenge.

Recent advances in mass spectrometry-based target deconvolution have enabled proteome-scale detection of ligand binding in a single experiment.^41^ Thermal proteome profiling assays and related approaches (*e.g.* the mass spectrometry-based cellular thermal shift assay (MS-CETSA) and the proteome integral solubility alteration assay, (PISA))^42,43^ are now staples in the label-free target identification toolkit. Despite widespread use, these assays suffer from drawbacks that hinder universal applicability. Live cell-based formats report on the downstream effects of compound treatment (*e.g.* changes in protein abundance) in addition to compound-induced thermal stabilization, which makes identification of direct protein targets more challenging.^44^ Lysate-based formats are useful in identifying direct targets but are performed under non-physiological conditions in which co-factors or co-substrates necessary for target engagement may become diluted or disrupted.^45,46^

To recapitulate intracellular binding dependencies, we reasoned that the addition of both ATP and the polypurine RNA required for rocaglate binding would promote target engagement in cell lysates. Here, we report an application of the PISA assay that allows for enhanced detection of rocaglate-induced clamping events. We show that ATP analogs and RNA substrates promote formation of DDX-RNA rocaglate receptor complexes in cell lysates, a requisite for rocaglate binding. We also show that limited proteolysis-mass spectrometry assays complement PISA to reveal new clamping targets. Structural docking-based analysis of DDX3X clamping by specific rocaglates serves as an example to inform structure-based design projects for five potential new additions to the rocaglate clamping spectrum. The approach described herein can be applied to screen for multicomponent interactions of other modalities and represents a new direction for stability proteomics as an evolving tool for drug discovery.

## EXPERIMENTAL PROCEDURES

### Cell culture

A549 cells (ATCC) were grown in Dulbecco’s Modified Eagle Medium supplemented with 10% fetal bovine serum and 1X penicillin-streptomycin solution. At ∼90% confluency, cells were washed with PBS, detached using TrypLE express, and washed 3X in HBSS. Cell pellets were flash frozen using liquid nitrogen and stored at -80 °C until use.

### Proteome integral solubility alteration (PISA) assay in cell lysates

Cell pellets were resuspended in ice-cold lysis buffer (20 mM HEPES, 138 mM NaCl, 5 mM KCl, 2 mM MgCl_2_, 2 mM CaCl_2_, cOmplete EDTA-free protease inhibitor, pH 7.4) and lysed via four successive freeze-thaw cycles in liquid nitrogen. Thawing was performed in a room temperature water bath. After the final freeze-thaw, lysates were clarified by centrifugation at 20,000 RCF and 4 °C for 15 minutes and transferred to a new LoBind tube. Lysate was assayed using a Pierce BCA Protein Assay Kit and adjusted to 2.0 mg/mL (2X experimental concentration).

Depending on the experiment, different 2X buffer solutions were prepared by supplementing the lysis buffer with 2X concentrations of an ATP analog and/or RNA probe – both of which were first made as stock solutions in LC-MS grade water – and a 2X concentration of the test compound or DMSO. Lysate was added 1:1 to the appropriate 2X buffer solution, gently mixed via pipette, and incubated at room temperature for 15 minutes. The final (1X) concentrations were as follows: ATP analog – 1 mM, RNA – 1 µM, test compounds – 10 µM, DMSO – 0.5%. After incubation, the 1X lysate was distributed across a 96-well PCR plate (20 *µ*L per well) and the plate was heated for 3 minutes in an Applied Biosystems ProFlex PCR System across a pre-defined temperature range consisting of six evenly spaced heating temperatures (50 to 64 °C for most experiments, 43 to 57 °C for **Figure S1D-E**). After heating, the PCR plate was placed on ice and an equal volume of lysate (18 *µ*L) was used to pool temperature points back to their original replicate samples. Samples were treated with 2 *µ*L of Benzonase (∼30 units) and incubated on ice for 30 minutes to digest nucleic acids. Protein aggregates were pelleted by centrifugation at 30,000 RCF and 4 °C for 30 minutes. 20 *µ*L of supernatant was transferred to new tubes containing 10 *µ*L of 3X sample preparation buffer (600 mM HEPES pH 8.5, 1.5% SDS, 30 mM TCEP, 120 mM chloroacetamide) and heated to 95 °C for 10 minutes to denature, reduce, and alkylate proteins. Samples were processed using single-pot solid phase-enhanced sample preparation (SP3).

### Limited proteolysis assay in cell lysates

Limited proteolysis was carried out following the single dose protocol as described in the publication, with minor deviation.^47^ Cell lysate and 2X experimental buffer were prepared as described above. Lysate and 2X buffer solution were mixed 1:1 and 55 *µ*L of 1X lysate was aliquoted across one row of a PCR plate. 5 *µ*L of proteinase K solution (0.2 *µ*g/*µ*L in water) was aliquoted across one row of the same PCR plate. The PCR plate was placed inside an Applied Biosystems ProFlex PCR System and equilibrated for 5 minutes at 25 °C. At the 5-minute mark, 50 *µ*L of lysate (50 *µ*g of protein) was transferred to the row containing 5 *µ*L of proteinase K using a 12-channel pipette. The proteinase K-lysate mixture was incubated for 4 minutes at 25 °C. At the 4-minute mark, the temperature program was automatically set to 99 °C and held for 10 minutes (with a 110 °C heated lid) to inactivate proteinase K. Samples were cooled to 4 °C and treated with Benzonase solution (11 *µ*L, ∼30 units) for 30 minutes to digest nucleic acids. Finally, 33 *µ*L of 3X sample preparation buffer (600 mM HEPES pH 8.5, 1.5% SDS, 30 mM TCEP, 120 mM chloroacetamide) was added to each sample, at which point the PCR plate was heated for 10 minutes at 95 °C for 10 minutes to resolubilize aggregates and reduce and alkylate proteins. After cooling, a volume of 20 *µ*L (∼10 *µ*g protein) was transferred to a new tube and processed using single-pot solid phase-enhanced sample preparation (SP3).

### Single-pot solid phase-enhanced sample preparation (SP3)

SP3 was carried out with minor deviation from the published protocol.^48,49^ Samples were mixed with ∼100 *µ*g of Sera-Mag carboxylate-modified magnetic beads (1:1, hydrophilic:hydrophobic beads) and proteins precipitated using acetonitrile. Samples were incubated on a ThermoMixer (24 °C, 1000 RPM) for 20 minutes, centrifuged for 1 minute at 500 RCF, and beads drawn to a magnet for 2 minutes. Beads were washed three times with 80% ethanol and once with 100% acetonitrile. Beads were air-dried, then resuspended in 50 *µ*L of digestion buffer (200 mM HEPES pH 8.5, 10 mM CaCl_2_, 0.01 *µ*g/*µ*L Trypsin/Lys-C mix), and proteins digested overnight at 37 °C with vortexing. After digestion, samples were reacted with ∼65 *µ*g of TMTpro multiplexing reagents for 1 hour. Samples were quenched with hydroxylamine, pooled, and evaporated to dryness in a SpeedVac. The pooled sample was resuspended in 2% acetonitrile, acidified with formic acid, and desalted using a Pierce Peptide Desalting Spin Column (ThermoFisher). The eluate was evaporated in a SpeedVac prior to high pH reversed-phase peptide fractionation.

### High pH reversed-phase peptide fractionation

TMTpro-labeled peptides were fractionated using high pH reversed-phase chromatography on an Agilent 1260 Infinity Capillary LC equipped with an XBridge Peptide BEH C18 column (300Å, 3.5*µ*m, 1mm x 150mm, Waters Corporation). Peptides were resuspended in 2% acetonitrile and 0.1% NH_4_OH, injected manually, and separated using a gradient of mobile phase A (2% acetonitrile, 0.1% NH_4_OH) to mobile phase B (98% acetonitrile, 0.1% NH_4_OH) at a flow rate of 75 *µ*L/min. Separation conditions were 100% mobile phase A for 5 minutes, a linear gradient to 12% mobile phase B for 5 minutes, a linear gradient to 32% mobile phase B for 35 minutes, a linear gradient to 45% mobile phase B for 8 minutes, a linear gradient to 70% mobile phase B for 1 minute, and a wash with 70% mobile phase B for 16 minutes (70 minutes total). Eluate fractions were collected from 20 to 68 minutes in 30 second time slices for a total of 96 fractions. Fractions were then concatenated into pooled fractions and evaporated in a SpeedVac. For PISA experiments, 12 fractions were analyzed^50^. For LiP-MS experiments, 24 fractions were analyzed.

### Mass spectrometry data acquisition

Fractions were separated on an Easy-nLC 1200 system connected to an Orbitrap Eclipse Tribrid mass spectrometer equipped with a FAIMS Pro Interface. Mobile phase A consisted of 2% acetonitrile, 0.1% formic acid in water and mobile phase B consisted of 0.1% formic acid in 80% acetonitrile. Peptides were first loaded onto an Acclaim PepMap C18 nano-trap column (100Å, 3*µ*m, 75*µ*m x 2cm, Thermo Scientific 164946) in mobile phase A, then resolved on an EASY-Spray column (100Å, 2*µ*m, 75*µ*m x 500mm, Thermo Scientific ES903). The flow rate was set to 250 nL/min. Peptides were separated using a 110 minute (LiP-MS) or 150 minute (PISA) method.

PISA data was collected using a real-time search SPS-MS3 method.^51^ The mass spectrometer was operated in positive ion mode with a spray voltage of 2500V and a capillary temperature of 275 °C. MS1 scans were acquired in the Orbitrap with a resolution of 60,000, scan range of 400-1600 m/z, AGC target set to standard, and a maximum injection time of 50 ms. For each FAIMS CV (-35, -50, -65V), the 10 most intense precursors were selected for MS2 scans in the ion trap. Additional filters included MIPS mode set to peptide, intensity threshold of 5e^3^, allowed charge states 2-6, and a 90 second dynamic exclusion window. MS2 scans were acquired in rapid mode with a 0.5 m/z isolation window, CID collision energy set to 35%, normalized AGC target of 200%, and maximum injection time set to 50 ms. MS2 spectra were searched in real-time against the Uniprot human reference proteome (UP000005640, downloaded February 8, 2023) combined with a list of common contaminants using the following settings: cysteine carbamidomethylation and TMTpro as fixed modifications, methionine oxidation (max. 2) as a variable modification, up to 2 missed cleavages, FDR filtering enabled, TMT SPS MS3 mode. Scoring thresholds applied to charge states of 2/3/4+ were Xcorr: 1/1/1, dCn: .05/.05/.05, and precursor PPM: 20/10/10. Successful searches triggered MS3 scans for TMTpro quantification. MS3 scans were acquired in the Orbitrap with a resolution of 50,000, HCD collision energy set to 55%, normalized AGC target of 350%, and maximum injection time of 200 ms.

LiP-MS data was collected using a TMTpro MS2 method. The mass spectrometer was operated in positive ion mode with a spray voltage of 2500V and a capillary temperature of 275 °C. Data dependent acquisition was performed with a cycle time of 1 second per FAIMS CV (-35, -50, -65V). MS1 scans were acquired in the Orbitrap with a resolution of 60,000, scan range of 400-1600 m/z, normalized AGC target of 100%, and maximum injection time set to automatic. Additional filters included MIPS mode set to peptide, intensity threshold of 2.5e^4^, allowed charge states 2-6, and a 60 second dynamic exclusion window. MS2 scans were acquired in the Orbitrap with a resolution of 50,000, HCD collision energy set to 35%, normalized AGC target of 200%, and maximum injection time set to automatic.

### Mass spectrometry data processing

Raw files from PISA experiments were searched and processed in Proteome Discoverer 3.1 using the standard RTS-SPS-MS3 consensus and processing workflow templates, which use both Comet and Sequest-HT to match identifications acquired during the real-time search. Spectra were searched against the Uniprot human reference proteome (UP000005640, downloaded February 8, 2023) with the following parameters: tryptic specificity, up to 2 missed cleavages allowed, peptide length: 7-30 aa, charge states: 2-6, precursor mass tolerance: 10 ppm, MS2 fragment mass tolerance: 0.6 Da, static modifications: cysteine carbamidomethylation and TMTpro on lysine and peptide N-terminus, variable modifications: methionine oxidation (max. 2). PSMs identified in Sequest-HT were rescored based on intensity using INFERYS. False discovery rates (target 0.01) were determined using Percolator with a target/decoy strategy^52^ in concatenated mode. Identifications were merged globally by search engine type and only high confidence peptides were used in subsequent steps. Protein quantification was performed with the following settings: unique peptides only, co-isolation threshold: 50%, minimum average reporter S/N threshold: 10, SPS mass matches threshold: 60%, all peptides used for protein roll-up. Intensity-based reporter abundances were normalized in subsequent steps (Proteomics data analysis section).

Raw files from LiP-MS experiments were converted to mzML format using MSConvert^53^ and processed in Fragpipe^54^ (v22.0) based on the parameters for the built-in TMT16 workflow. Spectra were searched against the Uniprot human reference proteome (UP000005640, downloaded February 8, 2023) appended with a list of common contaminants and decoys. The following parameters were used: semi-tryptic specificity, up to 2 missed cleavages allowed, peptide length: 7-50 aa, precursor mass tolerance: +/- 20 ppm, MS2 fragment mass tolerance: 20 ppm, fixed modifications: cysteine carbamidomethylation and TMTpro on lysine, variable modifications (max. 3): methionine oxidation, acetylation on protein N-termini, TMTpro on peptide N-termini. Peptide identifications were filtered at a 1% FDR. Peptide quantification was performed with the following settings: unique peptides only, minimum PSM probability: 0.9, minimum purity: 0.5, minimum summed reporter intensity (bottom 0.05). The abundance output file was grouped by peptide sequence, intensity values were log-transformed, and abundances were normalized in subsequent steps (Proteomics data analysis section).

### Proteomics data analysis

Additional data filtering and statistical analyses were conducted in R: A language and environment for Statistical Computing (R Foundation for Statistical Computing, http://www.R-project.org) using the Omics Notebook analysis pipeline^55^, which performs differential analysis using the limma R package.^56^

For PISA data, protein level results were exported from Proteome Discover. Contaminants, proteins with missing values, and proteins identified with <2 unique peptides were removed from the analysis. Abundance values were log-transformed and median normalized. For LiP-MS data, peptide level abundances were taken from the tmt-report output folder. Contaminants and peptides with missing values were removed from the analysis. Abundance values were log-transformed and quantile normalized. For all proteomics data sets, group comparisons were conducted using moderated t-tests with a Benjamini-Hochberg correction to contain the false discovery rate (5% for PISA, 1% for LiP-MS due to the higher number of statistical tests). Figures were built using the R package ggplot2 and GraphPad Prism (9.3.1).

### Purification of recombinant proteins

pET15b-His6-eIF4A1, pET15b-His6-eIF4A3, pColdII-His6-DDX3X, pET21a-His6-DDX21, or pColdII-His6-DDX50, were transformed into BL21 (DE3) codon+ *Escherichia coli* cells and cultured at 37 °C until the OD600 reached 0.6. Bacteria were induced with 1 mM IPTG and growth of cultures containing pET-backbone plasmids was continued for 3 h at 37 °C, while pColdII plasmid containing cultures were shifted to grow at 16 °C for 24h. Recombinant His6-eIF4A1, His6-eIF4A3, His6-DDX3X, His6-DDX21, and His6-DDX50 proteins were purified on a Ni^2+^-NTA agarose column followed by purification on a Q-Sepharose fast-flow matrix and elution with a linear salt gradient (100 to 500 mM KCl). Fractions containing protein were dialyzed against Buffer A (20 mM Tris-HCl [pH7.5], 10% glycerol, 0.1 mM ethylenediaminetetraacetic acid (EDTA)) overnight at 4 °C, and stored in aliquots at −80 °C.

### Fluorescence polarization assays

1.5 *μ*M recombinant eIF4A1, eIF4A3, DDX21, DDX50 or 7 *μ*M DDX3X protein was added to 10 nM FAM labeled poly (AG)_8_-RNA in binding buffer (14.4 mM HEPES-KOH [pH 8], 108 mM NaCl, 1 mM MgCl2, 14.4% glycerol, 0.1% DMSO, and 2 mM DTT) and 1 mM ATP in the presence or absence of the indicated compound (10 *μ*M). Following assembly, binding reactions were incubated for 10 min at room temperature (RT) in the dark, after which polarization values were determined using a Pherastar FS microplate reader (BMG Labtech). Data was analyzed using GraphPad Prism (9.3.1).

### Computational modeling

#### Ligand Preparation

Schrödinger’s LigPrep in the Maestro software environment (Version 14.1.138, Release 2024-3) was employed to prepare two compounds (Zotatifin and CR-1-31-B) for docking studies. During ligand preparation, possible protonation states prediction of the compounds was performed using Epik in the pH range of 7.4 +/- 2.0. For Zotatifin, LigPrep produced two predicted states for docking: the neutral species, and a protonated state where the dimethylamino nitrogen is positively charged. For CR-1-31-B, Epik produced only neutral species.

#### Rigid Receptor Docking

Glide docking was performed in Schrödinger’s Maestro software environment (Version 14.1.138, Release 2024-3). PDB X-ray structure 5ZC9 was prepared for use as the docking receptor through the default Protein Preparation Workflow, involving structure pre-processing, hydrogen-bond optimization, restrained minimization (S-OPLS force field, hydrogen atoms freely minimized and heavy atoms minimized to r.m.s.d. 0.3), and removal of water >4 Å from heteroatoms. For each compound, the highest-scored docking pose exhibiting the “canonical” rocaglate positioning observed in the 5ZC9 structure was selected for analysis.

#### Induced Fit Docking

Induced Fit Docking (IFD) was performed in Schrödinger’s Maestro software environment (Version 14.1.138, Release 2024-3). The ‘7LIU-C704A’ receptor was prepared from PDB X-ray structure 7LIU, point mutated to adenosine at DNA-RNA hybrid oligonucleotide residue C704 using pymol (Version 2.4.0, Schrödinger LLC), and was prepared for use as the docking receptor through the default Protein Preparation Workflow as described above. To define the binding site, the center-of-mass of each docked ligand was restrained to a 10 Å box positioned at the centroid of the following residues: Val328, Glu332, Gln360, Arg363 (from DDX3X), A704 and G705 (from the DNA-RNA hybrid oligonucleotide). During IFD, all protein side chains within 5 Å of the docked ligand were designated flexible, while all oligonucleotide side chains were designated as rigid. For each ligand, 11-14 ranked IFD poses were produced. For each compound, the highest-scored IFD pose exhibiting the “canonical” rocaglate positioning observed in the 5ZC9 structure was selected for analysis.

## RESULTS

### Thermal stabilization of pre-assembled rocaglate receptors

Given our goal of identifying novel interactions with DEAD-box (DDX) helicases, we set out to profile rocaglates with the lysate-based proteome integral solubility alternation (PISA) assay, a higher-throughput variant of thermal proteome profiling which detects compound-induced stability shifts (often described as altered solubility) in complex samples.^42,43^ Because lysate-based assays can be biased against co-factor- or substrate-dependent ligand binding^45,46^, we thought to promote clamping by pre-loading lysates with biochemical additives integral to the DDX helicase cycle.

Cycling DDXs undergo repeated rounds of RNA unwinding and release coupled with ATP hydrolysis.^57^ While rocaglate binding is not strictly ATP-dependent^31^, ATP promotes DDX-RNA interactions^58^ and therefore enhances the formation of rocaglate-competent DDX-RNA complexes **(Figure S1B)**. Moreover, Iwasaki and coworkers showed that RocA binds to eIF4A1-RNA complexes in the presence of ATP analogs that mimic each step of the hydrolytic cycle, including the post-ADP release state.^31^ Notably, clamping is most robust in the ATPase ground state mimicked using the non-hydrolysable ATP analog AMP-PNP **(Figure S1C)**. Finally, a large body of work supports the requirement for polypurine RNA in the rocaglate binding sites of eIF4A paralogs and DDX3X.^26,29,30,34^

Accordingly, we pre-loaded A549 lysates with AMP-PNP (1 mM) and a purine (AG)_8_ RNA 16mer (1 *µ*M), or water (control), and benchmarked PISA performance using the synthetic rocaglate CR-1-31-B **(Figure S1A)**, a confirmed binding partner of eIF4A paralogs and DDX3X with demonstrated biological activity.^2,13^ Treated lysates were distributed across a PCR plate, heated using a standard PISA thermal window (43 – 57 °C), then repooled and processed through a TMTpro workflow.

Results contrasted starkly between test conditions; no known targets were stabilized in the absence of additives, but eIF4A1, eIF4A3, and DDX3X were significantly stabilized in the presence of AMP-PNP and purine RNA **(Figures S1D-E)**. In addition, the Y-linked helicase DDX3Y – a speculative clamping target due to homology with DDX3X^29^ – was also stabilized, a finding afforded using a male derived cell line. Notably missing from the PISA profile, however, was eIF4A2 which, with a half-maximal melting temperature (T_m_) of ∼58 °C, fell victim to a key pitfall: reduced sensitivity for proteins with a high T_m_. Aiming to boost assay sensitivity, we looked to a key PISA study showing that higher thermal windows can magnify stabilization readouts **(Figure S1F)**.^59^ To guide selection of a more suitable thermal window, we pulled DDX thermal stability data from the Meltome Atlas^60^ and chose a range of 50 – 64 °C, which captures the bottom half of all DDX melt curves in the meltome data set **(Figure S1G)**.

We then evaluated the high thermal window using CR-1-31-B (10 *µ*M) in A549 lysates pre-loaded with either AMP-PNP (1 mM), a purine (AG)_8_ RNA 16mer (1 *µ*M), both additives, or neither (control). In line with results using a low thermal window, no DDXs were stabilized in the absence of additives **(Figure 1A)**, confirming that our initial negative result was not due to insufficient sensitivity; the use of only a single additive (AMP-PNP or purine RNA) likewise failed for identification of known targets **(Figures 1B-C)**. However, when both additives were included, all four bona-fide rocaglate targets were stabilized with markedly improved effect sizes **(Figures 1D-E)**. Beyond established clamping targets, DDX3Y was again stabilized, and DDX21 stabilization rose to significance, potentially adding another RNA helicase to the rocaglate clamping spectrum.

**Figure 1.**
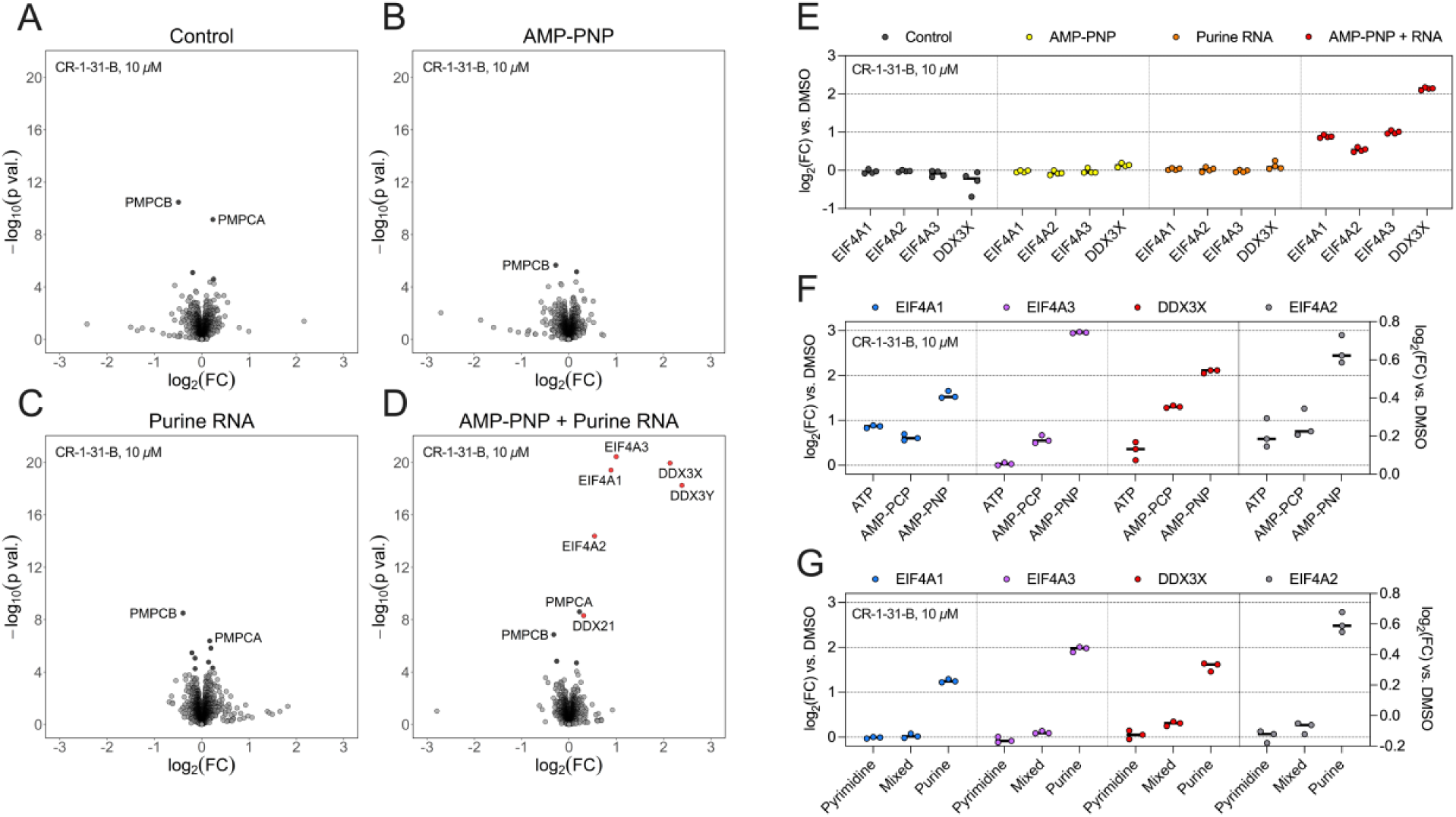
Additive helicase co-substrates promote thermal stabilization of rocaglate clamping targets. Volcano plots depicting PISA results for CR-1-31-B (10 *µ*M) in A549 lysate pre-loaded with **(A)** water (control), **(B)** AMP-PNP (1 mM), **(C)** purine (AG)_8_ RNA (1 *µ*M), or **(D)** both (n = 4). The x-axis represents fold change in stability relative to DMSO and the y-axis indicates the raw p value (BH adjusted p value < .05). Statistically significant proteins are colored black, statistically significant DDXs are colored red, and nonsignificant proteins are colored grey. **(E)** Extracted CR-1-31-B-induced fold changes in stability for established rocaglate clamping targets in the presence of water (control), AMP-PNP (1 mM), purine RNA (1 *µ*M), or both. Data corresponds to the volcano plots in Figures 1A through D. Lines indicate mean values. **(F)** Extracted CR-1-31-B-induced fold changes in stability for established rocaglate clamping targets in the presence of purine RNA (1 *µ*M) and the indicated ATP analog (1 mM) (n = 3). eIf4A1, eIF4A3, and DDX3X are plotted on the left y-axis and eIF4A2 is plotted on the right y-axis. Lines indicate mean values. **(G)** Extracted CR-1-31-B-induced fold changes in stability for established rocaglate clamping targets in the presence of AMP-PNP (1 mM) and a pyrimidine (CU)_8_ RNA 16mer (1 *µ*M), mixed (AU)_8_ RNA 16mer (1 *µ*M), or purine (AG)_8_ RNA 16mer (1 *µ*M) (n = 3). eIf4A1, eIF4A3, and DDX3X are plotted on the left y-axis and eIF4A2 is plotted on the right y-axis. Lines indicate mean values.

We verified that AMP-PNP enables more robust rocaglate-induced stabilization than the native ligand ATP and another non-hydrolyzable ATP analog, AMP-PCP **(Figure S1C)**, in a separate experiment using CR-1-31-B (10 *µ*M) and purine RNA (1 *µ*M). Indeed, while AMP-PCP did provide statistically significant stabilization for all four established clamping targets, effect sizes were substantially larger using AMP-PNP, whereas ATP only promoted stabilization of EIF4A1 **(Figure 1F)**, likely due to hydrolysis during the incubation. As expected, when we evaluated the importance of RNA base composition in another experiment using CR-1-31-B (10 *µ*M) and AMP-PNP (1 mM), all four *bona-fide* clamping targets were effectively unaltered in the presence of pyrimidine (CU)_8_ and mixed (AU)_8_ RNA 16mers, reflecting the purine specificity of the rocaglate targeting mechanism **(Figure 1G)**. Taken together, these results support a requirement for an ATP analog and exogenous purine RNA to promote the formation of stable rocaglate receptors in lysate-based PISA assays.

### PISA-based profiling of the rocaglate clamping spectrum

Having gained an understanding of rocaglate-specific PISA assay parameters, we forged ahead and evaluated a diverse set of rocaglates with varied core functionality. Our panel consisted of rocaglaol, RocA, silvestrol, CR-1-31-B, SDS-1-021^61,62^, and the clinical candidate Zotatifin (eFT226) **(Figure S1A)**. Each compound was tested at a concentration of 10 *µ*M in A549 cell lysate pre-loaded with AMP-PNP (1 mM) and purine (AG)_8_ RNA or pyrimidine (CU)_8_ RNA (1 *µ*M), totaling four separate experiments with each rocaglate-RNA pair tested in quadruplicate. We used the same batch of lysate across all four experiments to allow for head-to-head comparisons of fold-change (*i.e.* relative affinity). Full PISA profiles represented as volcano plots are provided in **Figures 2A** (purine RNA) and **S2** (pyrimidine RNA).

**Figure 2.**
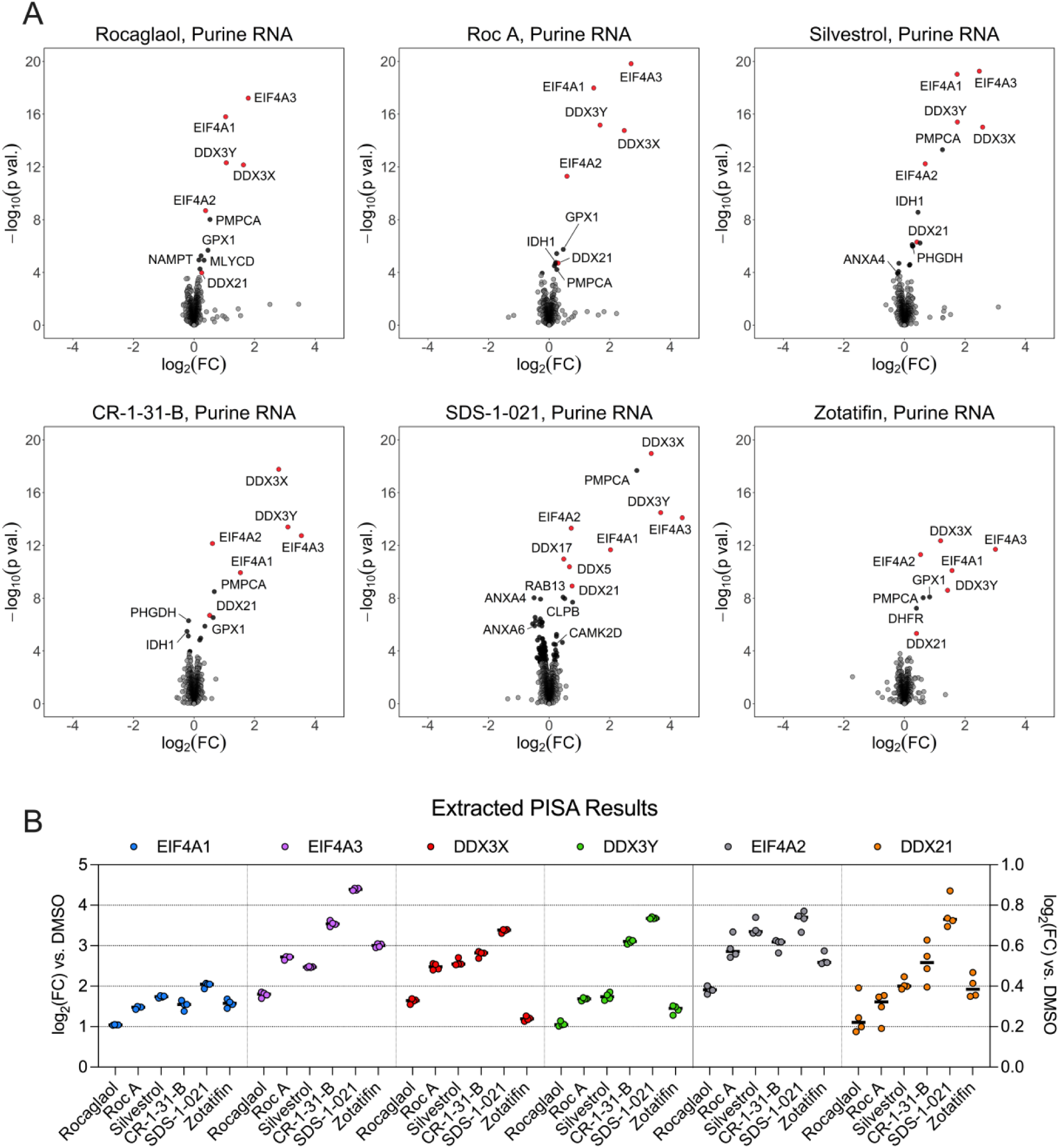
PISA-based profiling of the rocaglate clamping spectrum. **(A)** Volcano plots depicting PISA results for rocaglaol, Roc A, silvestrol, CR-1-31-B, SDS-1-021, and Zotatifin in A549 cell lysate pre-loaded with AMP-PNP (1 mM) and purine (AG)_8_ RNA (1 *µ*M) (n = 4). The x-axis represents fold change in stability relative to DMSO and the y-axis indicates the raw p value (BH adjusted p value < .05). Statistically significant proteins are colored black, statistically significant DDXs are colored red, and nonsignificant proteins are colored grey. **(B)** Extracted compound-induced fold changes in stability for commonly stabilized DDXs in the presence of purine RNA (1 *µ*M). eIf4A1, eIF4A3, DDX3X, and DDX3Y are plotted on the left y-axis; eIF4A2 and DDX21 are plotted on the right y-axis. Lines indicate mean values.

As anticipated, in the presence of purine RNA all rocaglates stabilized eIF4A1, eIF4A2, and, in line with the recent report from Naineni and coworkers, eIF4A3.^30^ Moreover, we found that each compound stabilized not only DDX3X but its close paralog DDX3Y. Zotatifin, touted for its eIF4A-selectivity, produced the smallest shift in DDX3 stabilization among the compounds tested. We note the broad dynamic range observed for stabilization of eIF4A1, eIF4A3, and DDX3 paralogs, which should allow for accurate comparison of relative affinities **(Figure 2B)**. In agreement with the results in **Figure 1G**, there was little notable eIF4A or DDX3 clamping when using pyrimidine RNA; the lone exception was for **SDS-1-021**, which weakly stabilized DDX3X and weakly destabilized eIF4A1 and eIF4A2 **(Figure S2)**.

Rocaglate-induced clamping of eIF4A and DDX3X is well understood, but whether rocaglates can similarly engage other members of the DDX family has remained an open question. In line with our findings in **Figure 1**, using purine RNA we found that all tested compounds significantly stabilize the helicase DDX21. SDS-1-021 also stabilized DDX21 in the presence of pyrimidine RNA, albeit to a lesser extent than with purine RNA, as well as DDX50, a close paralog of DDX21 with 55.6% sequence similarity.^63^ SDS-1-021 alone also stabilized the close paralogs DDX5 and DDX17, which overlap with 90% sequence similarity in their core domains.^64^ As for other putative targets, the use of a purine or pyrimidine probe had minimal impact on non-DDX PISA profiles **(Figures 2A and S2)**. For the majority of compounds, most differentially stabilized proteins between both RNA conditions were close enough to the statistical significance cutoff that their significance would reasonably fluctuate across experiments. Of note, however, in both RNA conditions we observed Zotatifin-induced stabilization of dihydrofolate reductase (DHFR), a well-characterized cancer and autoimmune disease target.^65^ Most importantly, by encompassing DDX5, DDX17, DDX21, DDX50, and paralogs of DDX3 and eIF4A, our PISA results potentially expand the rocaglate clamping spectrum from four to at least nine DDX proteins, all captured by the synthetic rocaglate congener SDS-1-021.

### Clamping validation with limited proteolysis and fluorescence polarization assays

Motivated by the potential of an expanded rocaglate clamping spectrum, we next used a complementary target identification method, limited proteolysis-mass spectrometry (LiP-MS), to validate some of our key PISA findings. LiP-MS is similar to the widely used drug affinity responsive target stability (DARTS) assay,^66^ which reports on ligand-induced changes to proteolytic cleavage patterns; DARTS traditionally employs immunoblot detection, whereas LiP-MS increases throughput and limits user bias *via* untargeted mass spectrometry detection.^47^ In principle, LiP-MS exploits ligand-induced conformational changes that differentially expose or shield a target protein from limited proteolysis by a non-specific protease. After a brief incubation, the non-specific protease is heat-inactivated, and samples are subjected to standard tryptic digestion. Compound and DMSO-treated samples are compared at the peptide level, and differentially quantified peptides implicate their parent proteins as putative targets.

We performed three independent LiP-MS experiments with a subset of synthetic rocaglates: CR-1-31-B, SDS-1-021, and Zotatifin (10 *µ*M each). Based on our PISA screens, we tested each compound in A549 lysate pre-loaded with AMP-PNP (1 mM) and (AG)_8_ RNA (1 *µ*M). After an equilibrating incubation with the test compound or DMSO, samples were subjected to limited proteolysis with proteinase K, followed by heat inactivation. We processed the samples through TMTpro workflows and performed differential analysis using both tryptic and semi-tryptic peptide abundance values. Representative volcano plots for each experiment are provided in **Figures 3A-C**. A complete list of all differentially cleaved peptides can be found in the Supplementary Data Tables.

**Figure 3.**
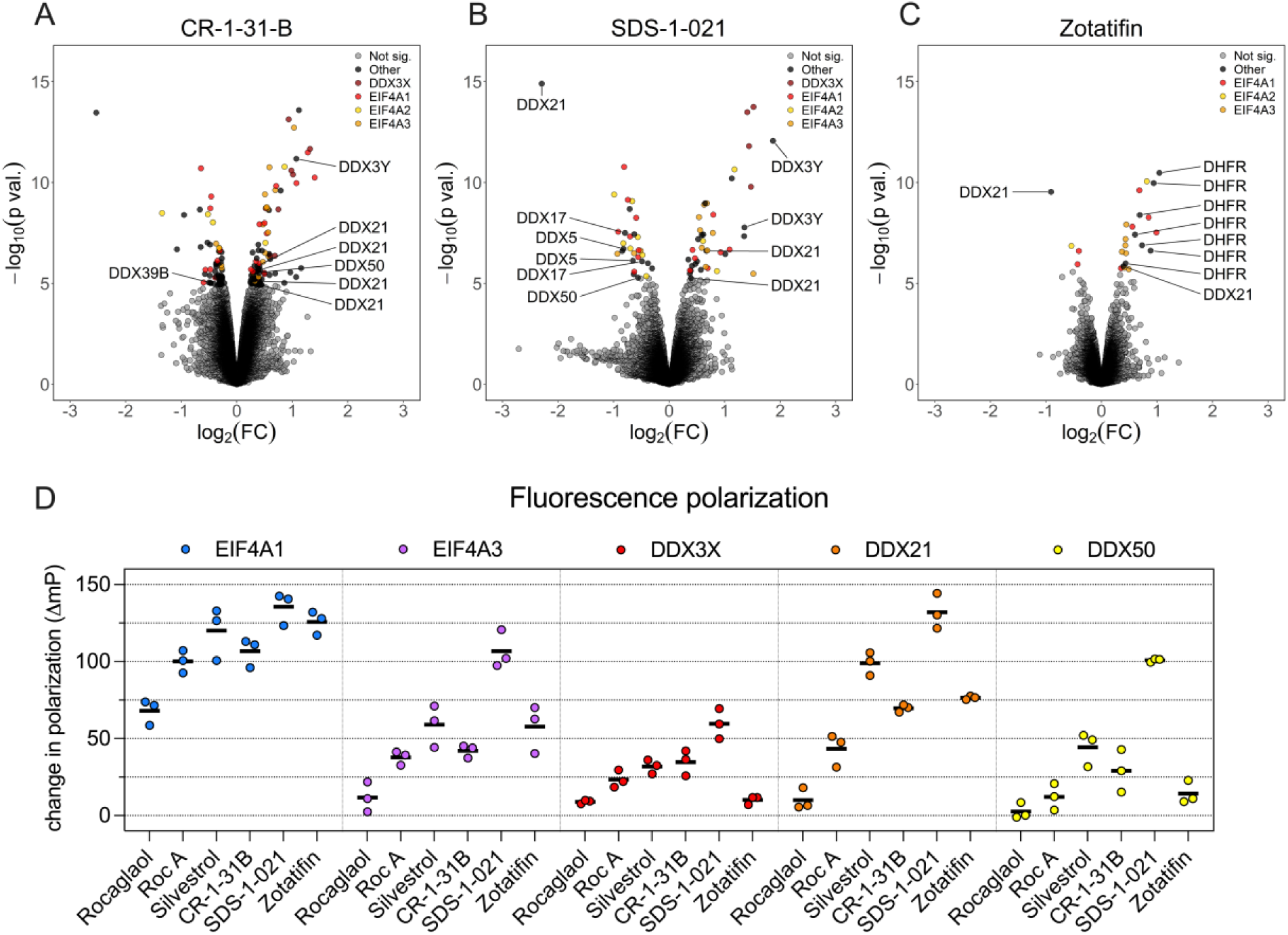
Clamping validation with limited proteolysis and fluorescence polarization assays. Volcano plots depicting limited proteolysis results for three independent experiments testing **(A)** CR-1-31-B, **(B)** SDS-1-021, or **(C)** Zotatifin at 10 *µ*M in A549 cell lysate pre-loaded with AMP-PNP (1 mM) and purine (AG)_8_ RNA (1 *µ*M) (n = 6). Plot points are labeled with protein names corresponding to unique tryptic or semi-tryptic peptides that were differentially cleaved between compound and DMSO treated samples. The x-axis represents fold change in peptide abundance relative to DMSO and the y-axis indicates the raw p value (BH adjusted p value < .01). Unique peptides stemming from eIF4A paralogs or DDX3X are colored as indicated and not labeled; unique peptides stemming from other significant DDXs are colored black and labeled by protein; nonsignificant proteins are colored grey. **(D)** Recombinant DDX proteins were incubated with a 5’-FAM-labeled (AG)_8_ RNA probe, then treated with test compound or DMSO (n = 3). Polarized fluorescence emission was measured in a polarimeter, and the DMSO-treated baseline was subtracted from compound-treated polarization to derive the compound-induced change in fluorescence polarization (ΔmP). Lines indicate mean values.

As expected, all three compounds altered the proteolytic profiles of eIF4A1, eIF4A2, and eIF4A3 which resulted in multiple differentially abundant peptides for each paralog. DDX3X and DDX3Y peptides were among the most affected by CR-1-31-B and SDS-1-021, but not Zotatifin, consistent with our PISA results showing Zotatifin as the weakest DDX3 stabilizer among the synthetic rocaglates **(Figure 2B)**. As for alternative DDX clamping candidates, in CR-1-31-B-treated samples we observed four differentially abundant peptides from DDX21, one from DDX50, and one from DDX39B though the effect sizes were small **(Figure 3A)**. Results for SDS-1-021 mostly recapitulated our PISA findings – we detected changes to three peptides from DDX21, one from DDX50, and two peptides each from DDX5 and DDX17, with larger effect sizes than seen for CR-1-31-B **(Figure 3B)**. Notably, Zotatifin treatment resulted in two differentially abundant peptides from DDX21 **(Figure 3C)**. Strikingly, of the 25 differentially abundant peptides detected in the Zotatifin experiment, seven belonged to DHFR. Nevertheless, in conjunction with our PISA data, these LiP-MS results further implicate DDX3Y, DDX5, DDX17, DDX21, and DDX50 as DEAD-box helicase leads for expansion of the rocaglate clamping spectrum.

To firmly establish additional DDXs as direct rocaglate targets, we employed a previously developed fluorescence polarization (FP) assay using a 5’-fluorescein amidite (5’-FAM)-labeled (AG)_8_ RNA probe.^26^ After successful production of recombinant eIF4A1, eIF4A3, DDX3X, DDX21, and DDX50, we used FP to evaluate our panel of rocaglates for their ability to clamp these proteins to labeled polypurine RNA, reported as a change in polarized fluorescence (ΔmP) relative to DMSO-treated controls. Gratifyingly, DDX21 and DDX50 clamped purine RNA to some extent in the presence of most compounds **(Figure 3D)**. This effect was the most profound for SDS-1-021 which was the strongest clamper for all proteins tested. Results for Zotatifin largely echoed PISA and LiP-MS data, with moderate clamping of DDX21 and weak clamping of DDX3X observed. We also note that our FP-derived clamping values – a proxy for relative affinity – generally align with our PISA-derived stabilization values **(Figure 2B)**, supporting the use of PISA as a tool to gauge relative affinity and selectivity. These results establish DDX21 and DDX50 as tractable DEAD-box helicase targets for future rocaglate medicinal chemistry campaigns.

### Computational modeling of rocaglate binding to DDX proteins

The heterogeneity of target profiles observed across the tested compounds raises the intriguing possibility to design rocaglates that specifically target members of the DDX family. To that end, we wanted to understand the molecular basis for Zotatifin’s apparent targeting away from DDX3X in comparison to other synthetic rocaglates. Zotatifin is structurally similar to CR-1-31-B **(Figure 3A)**, with a nitrogen replacing C7 of the A-ring, a nitrile at the C4’ position of the B-ring, and an N,N-dimethylaminomethylene substituent at the C2-position. We first used conventional Glide docking^67,68^ to model both CR-1-31-B and Zotatifin into their shared target, eIF4A1, using a published RocA-(AG)_5_-eIF4A1 X-ray crystal structure (PDB ID: 5ZC9). Both docking exercises showed the respective ligands bound in a comparable position to RocA (hereafter referred to as the “canonical” rocaglate pose), with tightly sandwiched π-stacking interactions between the A- and B-rings and (AG)_5_ RNA **(Figures 4B-C and S3A-B)**. Beyond ligand-RNA interactions, RocA has been shown to engage in limited contacts with eIF4A1 itself, the two most prominent being a hydrogen bond between the RocA C2 amide carbonyl and eIF4A1 Gln195, and a parallel-displaced π-stacking interaction between the RocA C-ring and eIF4A1 Phe163.^34^ Both interactions were fully recapitulated in the top eIF4A1 docking pose for CR-1-31-B (Glide Gscore: -8.257 kcal/mol) **(Figures 4B and S3A)**. For the top Zotatifin-eIF4A1 pose (Glide Gscore: -8.144 kcal/mol) **(Figures 4C and S3B)**, the expected π-stacking to Phe163 was also observed. Interestingly, while Zotatifin lacks a carbonyl at the C2 position, docking predicted the lone pair of the C2 dimethylamino nitrogen to sit 2.2 Å from the nearest Gln195 sidechain proton, indicative of a strong hydrogen bond.

**Figure 4.**
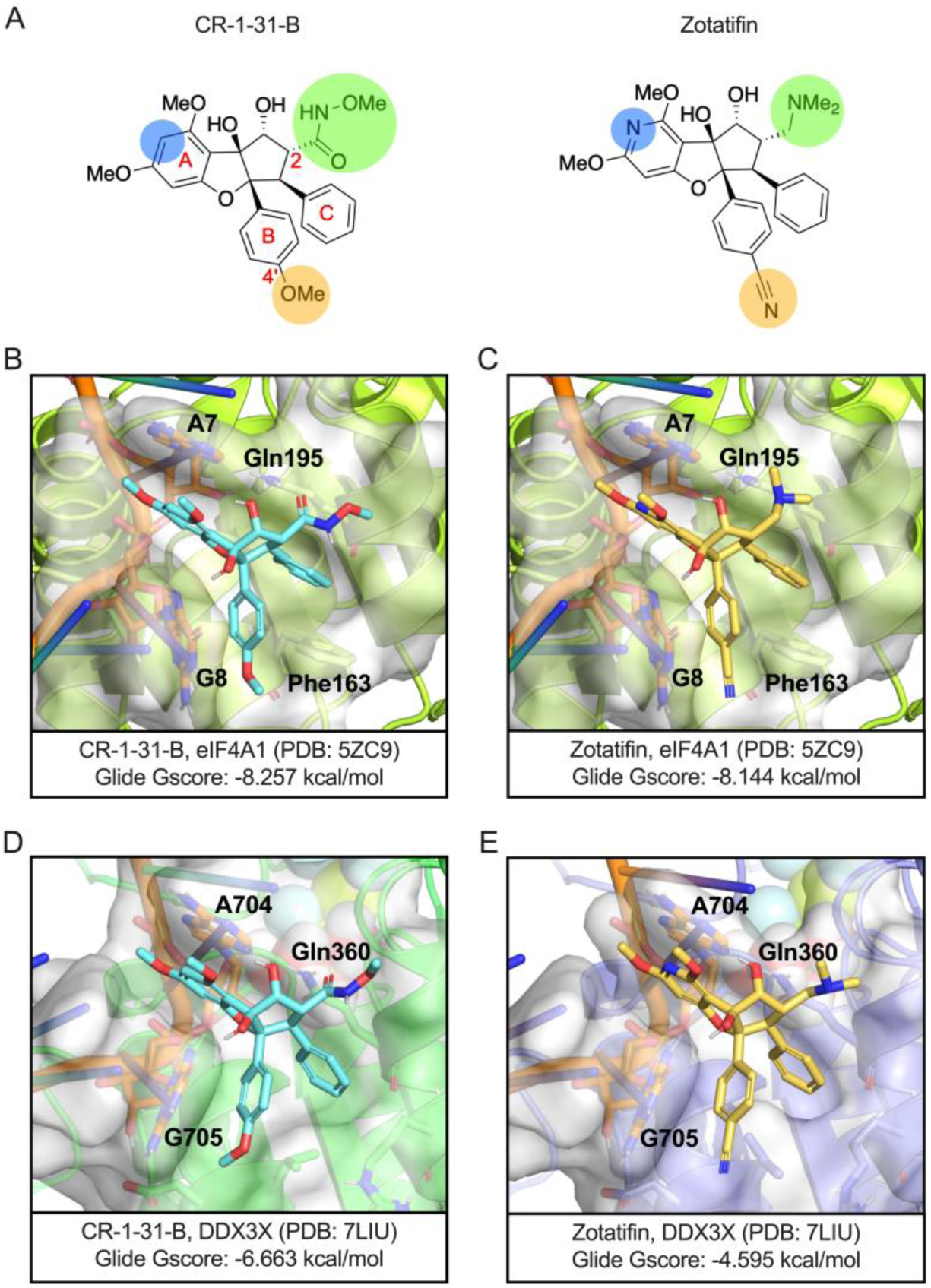
Docking of synthetic rocaglates into eIF4A1 and DDX3X. Rocaglate binding was analyzed *via* computational docking of CR1-31-B and Zotatifin into X-ray crystal structures for eIF4A1 bound to (AG)_5_ RNA (PDB: 5ZC9), and DDX3X bound to an RNA-DNA hybrid (PDB: 7LIU). Note that single letter codes (*e.g.* A7) refer to RNA bases and three letter codes refer to amino acid residues. **(A)** Chemical structures of CR-1-31-B and Zotatifin with key structural differences highlighted. **(B)** Pose for CR-1-31-B bound to an eIF4A1-RNA structure (PDB: 5ZC9) obtained from rigid-receptor docking. **(C)** Pose for Zotatifin bound to an eIF4A1-RNA structure (PDB: 5ZC9) obtained from rigid-receptor docking. **(D)** Pose for CR-1-31-B bound to a DDX3X-RNA/DNA structure (PDB: 7LIU) obtained from Induced Fit Docking. **(E)** Pose for Zotatifin bound to a DDX3X-RNA/DNA structure (PDB: 7LIU) obtained from Induced Fit Docking.

To better understand the molecular basis for their differential targeting, we next attempted to model the two ligands into DDX3X. Here, we used an X-ray structure of DDX3X in a “rocaglate-competent” post-unwound configuration (PDB: 7LIU), in which the helicase is bound to a DNA:RNA hybrid. Unfortunately, conventional (flexible-ligand/rigid-receptor) Glide docking failed to produce the canonical rocaglate pose for either CR-1-31-B or Zotatifin into this structure. Consequently, we opted to deploy Schrödinger’s flexible-ligand/flexible receptor Induced Fit Docking (IFD) workflow^69^ given that receptors are well-precedented to undergo subtle conformational changes to accommodate ligand binding.^13,70^ IFD modeling, which maintains a rigid protein backbone while allowing flexibility of sidechain residues, allows for structural adjustments at the putative rocaglate binding site that may serve to both avoid steric clashes and access new favorable receptor-ligand binding interactions.

Gratifyingly, using a rocaglate competent model, where a single cytosine-to-adenine point mutation was performed on the RNA:DNA hybrid (7LIU-C704A), the top-scored IFD poses for neutral CR-1-31-B **(Figures 4D and S3C)** and Zotatifin **(Figure 4E)** docked into DDX3X both achieved the canonical rocaglate pose, albeit with different predicted binding affinities (Glide scores: -6.663 kcal/mol for CR-1-31-B *vs*. -4.595 kcal/mol for Zotatifin). For both ligands, IFD predicted π-stacking interactions between the rocaglate A- and B-rings to A704 and G705 of the RNA:DNA hybrid, respectively. The key C-ring binding residue in eIF4A1, Phe163, corresponds to residue Val328 in DDX3X **(Figure S3C)**, precluding any possibility of π-stacking for either compound. In the case of CR-1-31-B, we observed a hydrogen bond between the carbonyl group of the C2-hydroxamic ester and the sidechain of DDX3X Gln360 **(Figures 4D and S3C)**, a residue analogous to eIF4A1’s Gln195 **(Figure S3A)**. In contrast, for Zotatifin no interactions were observed with Gln360. Due to positional differences for the ligand core across the two poses **(Figure S3D)**, in the Zotatifin-DDX3X IFD structure the C2-pendant amine is turned away from Gln360, eliminating any potential to form a hydrogen bond with this key rocaglate-binding residue.^29^ We postulate that this difference in ability to bind Gln360 is a key driver of the differential targeting of DDX3X by the two ligands. Taken together, these modeling experiments provide a potential structural basis for the experimentally observed differential targeting of DDX3X by CR-1-31-B and Zotatifin.

## DISCUSSION

To date, the rocaglate clamping spectrum has been established for four DDX proteins: eIF4A1, eIF4A2, eIF4A3, and DDX3X. Here, we describe a stability proteomics approach that enables detection of molecular clamping at proteomic scale. Using PISA and LiP-MS, we provide evidence of an expanded rocaglate clamping spectrum, with validation for DDX21 and DDX50 engagement by multiple widely used rocaglates, and implication of DDX3Y, DDX5, and DDX17 as rocaglate-ligandable RNA helicases. We show that these findings are conditioned upon target class-specific assay conditions, including additives that promote *in vitro* assembly of rocaglate receptor complexes. Our data also suggests DHFR as a potential off-target of Zotatifin, providing a starting point to mechanistically probe toxicity or polypharmacology. In keeping with our overarching goal of structure-based rocaglate design, we propose a molecular basis to explain differences in DDX3X stabilization (*i.e.* affinity) through comparative docking of Zotatifin and CR-1-31-B. More broadly, our study provides a generalizable approach for chemical proteomic stability screens of other interfacial ligands, including pateamines, camptothecins, or molecular glues.

Our study is not the first chemical proteomics-based survey for rocaglate targets. It does, however, demonstrate advantages over prior methods which relied on affinity enrichment^71,72^ or covalent labeling.^29^ For example, Chambers’ pull-down of a biotinylated silvestrol probe did not employ additive RNA or ATP analogs, a point that Chen notes.^29^ In this case, DDXs of low affinity and abundance likely fell victim to washing in the absence of these ectopic reagents. Chen’s approach, which did include RNA and AMP-PNP, used an O-nitrobenzoxadiazole *(O-*NBD)-based RocA probe to label proximal lysines, which are not always situated near binding sites. In both examples, rocaglate derivatization has the potential to alter target binding profiles and yield spurious results. In contrast, PISA and LiP-MS require no modification to the active compound, allowing for more efficient use of synthetic chemistry resources. On the other hand, stability proteomics methods have a degree of uncertainty as to whether the relevant target is sufficiently altered under the chosen perturbation conditions (*e.g.* dose, heat, proteolysis). Only using a higher thermal window were we able to detect EIF4A2 and DDX21 clamping **(Figures 1D and S1E);** however, higher thermal windows come with the cost of reduced coverage as a larger fraction of the proteome is melted away. For these reasons, multiple complementary methods are advisable in target identification campaigns.

Why does AMP-PNP offer improved assay performance over the native ligand ATP? Structural studies of ATPases often employ non-hydrolysable ATP analogs to trap enzymes in a pre-hydrolytic state.^73^ Others have shown that DDX-RNA complexes dissociate rapidly, with half-lives of seconds to minutes in the presence of ATP, whereas AMP-PNP-bound complexes can persist for hours.^31^ The labile nature of ATP (*i.e.* hydrolysis) likely manifests as a suboptimal distribution of cycling DDXs in ATP-treated lysates, given that purine-bound DDXs are the most suitable rocaglate receptors. Whether AMP-PNP is the most useful ATP analog for investigations with different RNA additives remains a topic for future investigations. We speculate that experiments with more complex RNA molecules (*i.e.* structured RNA)^74^ may benefit from the use of ATP, as it could allow DDX-RNA complexes to cycle until a rocaglate-competent binding site is formed. Investigations around the nucleic acid structural preferences of different DDX proteins may shed light on biological processes and disease states ripe for rocaglate targeting.

Regarding novel DDX proteins, we establish DDX21 and DDX50 as *bona-fide* rocaglate targets amenable to RNA clamping. DDX21 has reported roles in ribosome biogenesis, innate immunity, and glucose sensing, among several others.^37,75^ Its close paralog DDX50 was recently characterized as a viral sensor involved in Dengue virus infection.^63^ Their sequence similarity may suggest parallel targeting as with eIF4A paralogs, though closer study is warranted at this time. Thus far, we have not had the opportunity to confirm DDX5 and DDX17 engagement using recombinant protein, but their high degree of core domain similarity may suggest a parallel targeting effect as well.

We also note that Zotatifin, an eIF4A-selective rocaglate that appears to have been successfully tuned away from DDX3, also altered DHFR and DDX21 in both PISA and LiP-MS assays. DHFR, a historically well-studied cancer and auto-immune target, is inhibited by folate antagonists (*e.g.* methotrexate), which have been shown to also possess *in vitro* antiviral activity.^65,76–78^ Given Zotatifin’s status as a clinical candidate for both ER+ breast cancer and SARS-CoV-2 infection, our findings warrant follow-up to determine whether DHFR and DDX21 contribute to Zotatifin’s efficacy or pose liabilities.

Zotatifin was optimized for eIF4A inhibition in a campaign that also targeted improved drug-like properties through reduced lipophilicity and increased aqueous solubility, which was presumably a key motivation behind installing the C2 tertiary amine, C4’ nitrile, and nitrogenated A-ring. However, our modeling and experimental data suggest that these modifications also provided a ligand relatively selective for eIF4A. We posit that for Zotatifin, an *in silico*-predicted loss of Gln360 H-bonding capability at C2 significantly hampers DDX3X engagement. In the case of CR-1-31-B, however, our modeling predicts a retained interaction with Gln360 that appears to offset the loss of π-π stacking arising from the Phe→Val mutation, a phenomenon that we expect is extendable to other C2 carbonylated DDX3X-targeting rocaglates such as silvestrol. While not directly interrogated by our modeling, we note it is also possible that subtle electronic differences in the pyridine A-ring of Zotatifin compared with phenyl A-ring of CR-1-31-B may impact the π-stacking interactions with RNA in the ternary complex.^7^ While the predicted conformational changes must still be validated experimentally, this modeling study nonetheless introduces the potential to strategically tune rocaglates toward or away from specific helicases. Currently, the main barrier to this goal is a paucity of useful data to inform structure-based drug design. Using PISA and allied techniques, we hope to catalyze rocaglate drug discovery by shifting medicinal chemistry focus toward actionable DDX targets, opening the door for next generation of RNA-protein molecular clamps.

## ASSOCIATED CONTENT

### Data availability statement

The mass spectrometry proteomics data have been deposited to the ProteomeXchange Consortium via the PRIDE ^79^ partner repository with the dataset identifier PXD058802.

### Supporting information

The Supporting Information is available free of charge at [URL].

- Supplementary figures as referenced in the text (SupplementaryFigures.pdf)
- Supplementary data tables with processed statistical results for PISA experiments (SupplementaryData1.xlsx) and LiP-MS experiments (SupplementaryData2.xlsx)

## AUTHOR INFORMATION

### Author Contributions

SIG and JAP designed and supervised the study. SIG and SBG performed the PISA experiments. SIG and ACF performed the LiP-MS experiments. SKN and RC performed the fluorescence polarization experiments. SKN, RC, and KP generated the purified recombinant proteins. SH supervised the fluorescence polarization experiments. ZW and LEB performed and supervised the docking experiments. AE supervised the mass spectrometry experiments. SIG, ZW, LEB, and JAP wrote the manuscript with inputs from all other authors.

## ACKNOWLEDGEMENTS

We dedicate this article to the memory of Professor Jerry Pelletier, a pioneer in the field of RNA biology whose work laid the foundation for rocaglate drug discovery. We thank the National Institutes of Health (R35 GM 118173) for financial support and for support in funding an ultra-high precision mass spectrometer (S10OD026807). We also thank the National Institute of General Medical Sciences (T32 GM 008541) and the Canadian Institute of Health Research (PJT-186015, PJT-183610) for additional funding. We thank Michelle Nguyen and Dr. John Connor (BU) for providing A549 cells. CETSA ® (*i.e.* PISA) data was generated under an agreement with Pelago Bioscience AB who own the CETSA patent family. Computational work at Boston University reported in this paper was performed on the Shared Computing Cluster (SCC) which is administered by Boston University’s Research Computing Services.

