## Supplementary Figures for "Discovery of RNA-Protein Molecular Clamps Using Proteome-Wide Stability Assays"

**Using Proteome-Wide Stability Assays**

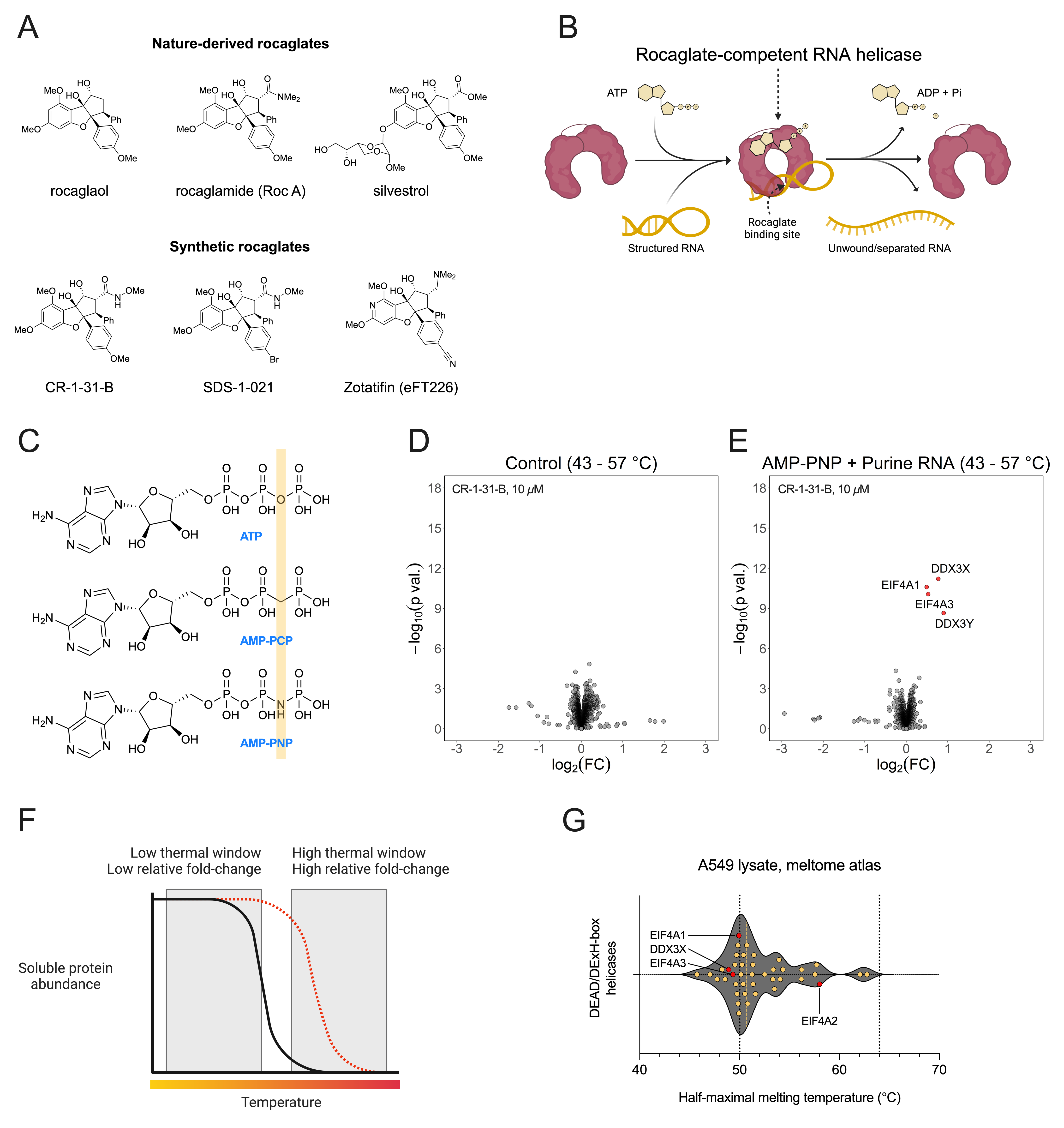

**Figure S1. Thermal stabilization of pre-assembled rocaglate receptors. (A)** Rocaglates used in this study. **(B)** ATP-dependent RNA binding by a DEAD-box helicase. ATP and RNA binding are coupled, as are ATP hydrolysis and RNA release. **(C)** ATP analogs used in this study with differences highlighted. **(D)** PISA results for CR-1-31-B (10 *µ*M) in A549 lysate without additives using a standard thermal window of 43 – 57 °C (n = 4). **(E)** PISA results for CR-1-31-B (10 *µ*M) in A549 lysate pre-loaded with AMP-PNP (1 mM) and (AG)_8_ purine RNA (1 *µ*M) using a standard thermal window of 43 – 57 °C (n = 4). **(F)** Performing PISA with high thermal windows leads to higher fold changes than using low thermal windows. Thermal windows are indicated by the grey boxes. **(G)** Distribution of DDX protein T_m_ values in A549 cell lysate taken from Meltome Atlas.

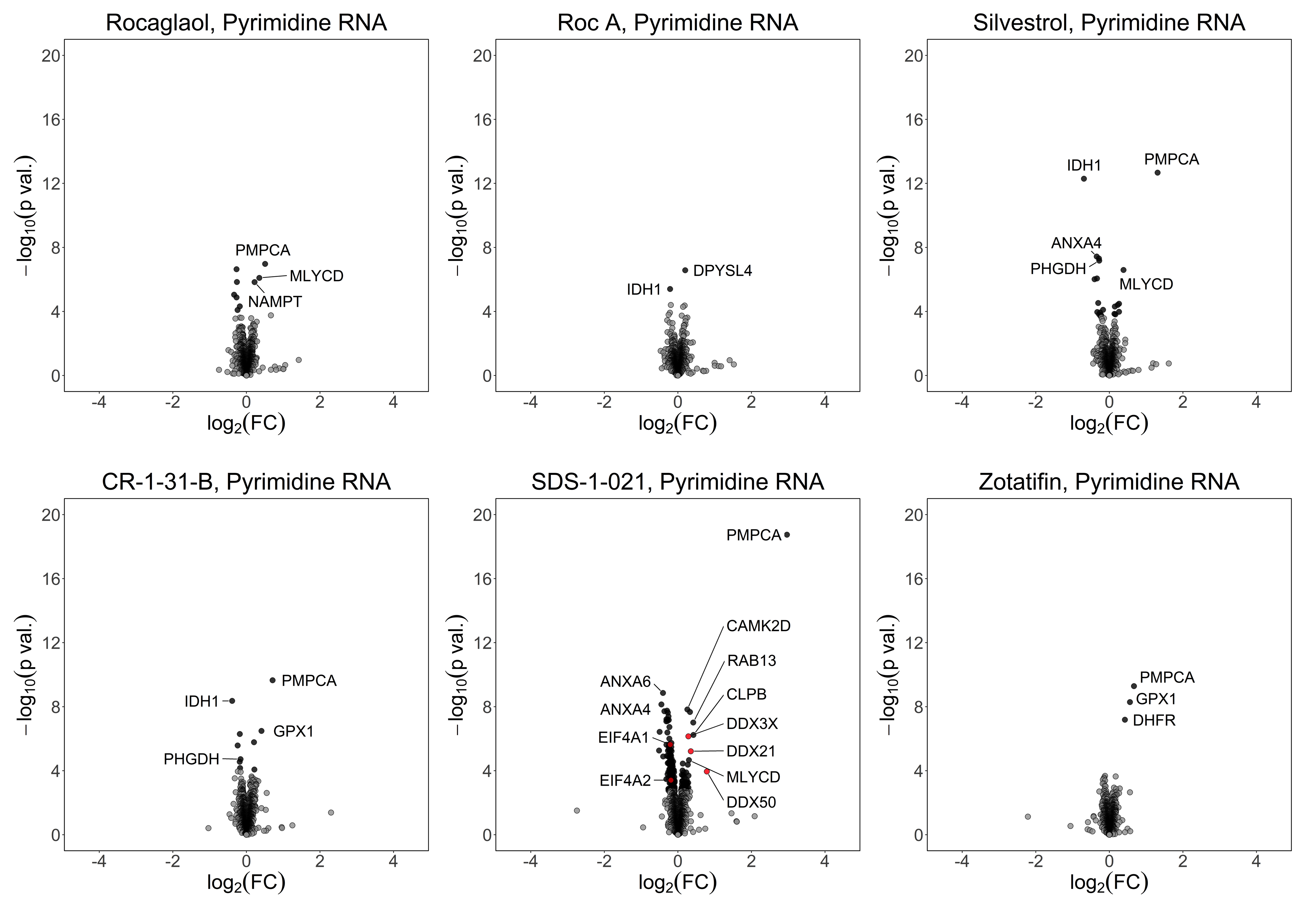

**Figure S2. PISA profiling with a panel of rocaglates using pyrimidine RNA.** Volcano plots depicting PISA results for rocaglaol, Roc A, silvestrol, CR-1-31-B, SDS-1-021, and Zotatifin in A549 cell lysate pre-loaded with AMP-PNP (1 mM) and pyrimidine (CU)_8_ RNA (1 *µ*M) (n = 4). The x-axis represents fold change in stability relative to DMSO and the y-axis indicates the raw p value (BH adjusted p value < .05). Statistically significant proteins are colored black, statistically significant DDXs are colored red, and nonsignificant proteins are colored grey.

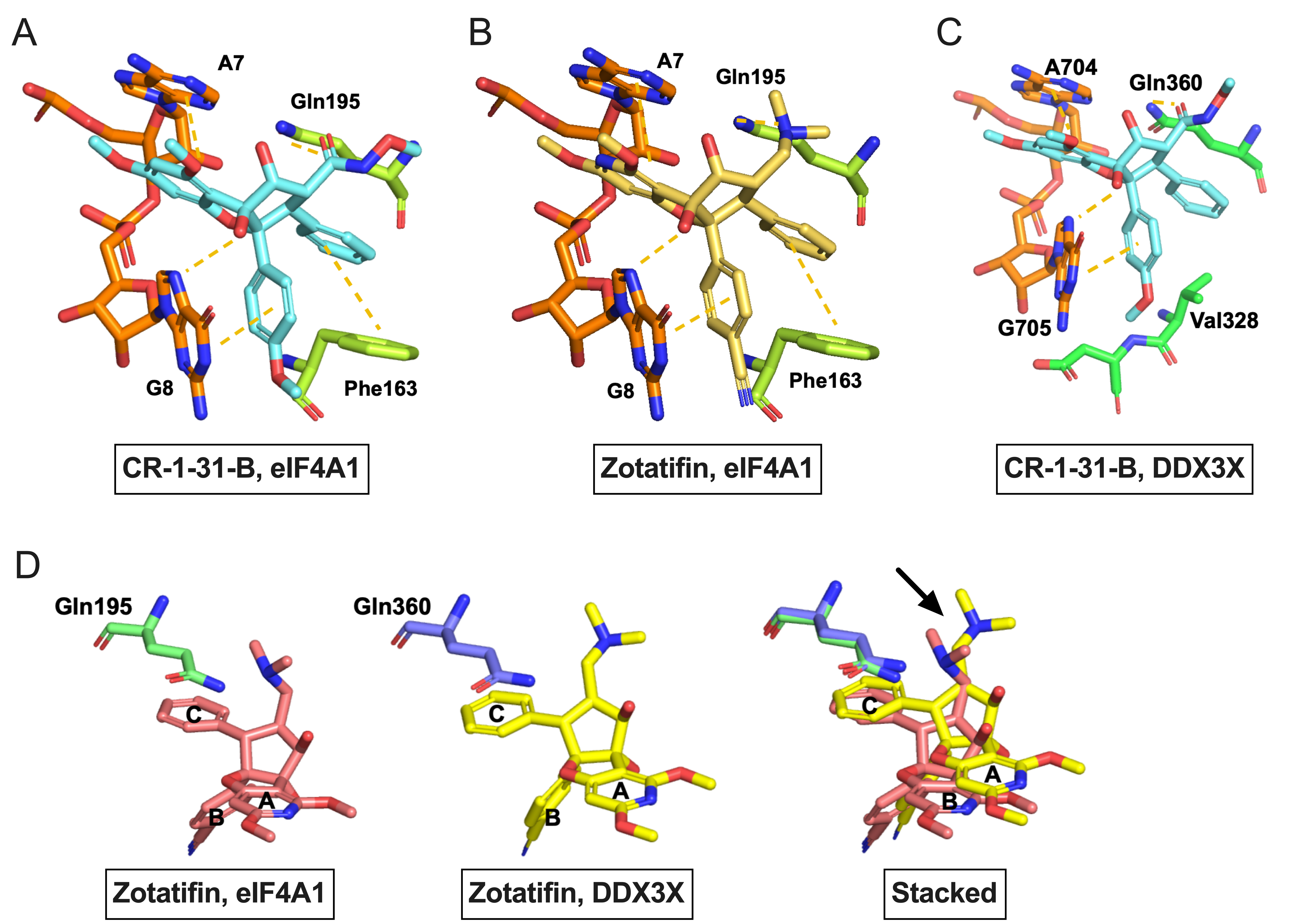

**Figure S3. Docking of synthetic rocaglates into eIF4A1 and DDX3X.** Poses for CR-1-31-B **(A)** and Zotatifin **(B)** docked into a published RocA-(AG)_5_-eIF4A1 X-ray crystal structure (PDB ID: 5ZC9) with key binding interactions indicated by the dotted lines. Note that single letter codes (*e.g.* A7) refer to RNA bases and three letter codes refer to amino acid residues. **(C)** Induced fit docking (IFD) of CR-1-31-B into a DDX3X-RNA/DNA hybrid structure (PDB ID: 7LIU) with key binding interactions indicated by the dotted lines. **(D)** An overlay of modeling results for Zotatifin docked into eIF4A1 (PDB ID: 5ZC9) and DDX3X (PDB ID: 7LIU). Arrow points to the pendant amine turned away from Gln360 in the induced fit model.
